# The genetic architecture underlying body-size traits plasticity over different temperatures and developmental stages in *Caenorhabditis elegans*

**DOI:** 10.1101/2021.04.21.440777

**Authors:** Muhammad I. Maulana, Joost A.G. Riksen, Basten L. Snoek, Jan E. Kammenga, Mark G. Sterken

## Abstract

Most ectotherms obey the temperature-size rule, meaning they grow larger in a colder environment. This raises the question of how the interplay between genes and temperature affect the body size of ectotherms. Despite the growing body of literature on the physiological life-history and molecular genetic mechanism underlying the temperature-size rule, the overall genetic architecture orchestrating this complex phenotype is not yet fully understood. One approach to identify genetic regulators of complex phenotypes is Quantitative Trait Locus (QTL) mapping. Here, we explore the genetic architecture of body size phenotypes, and plasticity of body-size phenotypes in different temperatures using *Caenorhabditis elegans* as a model ectotherm. We used 40 recombinant inbred lines (RILs) derived from N2 and CB4856, which were reared at four different temperatures (16°C, 20°C, 24°C, and 26°C) and measured at two developmental stages (L4 and adult). The animals were measured for body length, width at vulva, body volume, length/width ratio, and seven other body-size traits. The genetically diverse RILs varied in their body-size phenotypes with heritabilities ranging from 0.0 to 0.99. We detected 18 QTL underlying the body-size traits across all treatment combinations, with the majority clustering on Chromosome X. We hypothesize that the Chromosome X QTL could result from a known pleiotropic regulator – *npr-1 –* known to affect the body size of *C. elegans* through behavioral changes. We also found five plasticity QTL of body-size which three of them colocalized with some body-size QTL at certain temperature. In conclusion, our findings shed more light on multiple loci affecting body size plasticity and the possibility of co-regulation of traits and traits plasticity by the same loci under different environment.

## Introduction

The body size of ectotherms such as invertebrates, insects and fish are negatively correlated with their ambient temperature, where warmer environments result in smaller body-size. Besides body-size, the ectotherms’ life-history traits are also strongly affected by temperatures. Phenotypic plasticity (the phenotypes that can be expressed by a single genotype at different environmental conditions) due to temperature changes has been studied widely for many different ectotherms, including evolutionary, ecological, physiological, and molecular investigations (Beldade et al., 2011; Callahan et al., 2005; Lafuente & Beldade, 2019; Scheiner, 1993; Via et al., 1995).

In particular, body size plasticity has been studied well, aiming to understand why ectotherms grow larger at lower temperatures, a process called the temperature-size rule (Angilletta & Dunham, 2003; Atkinson, 1994; Ghosh et al., 2013; Van Voorhies, 1996). Atkinson, 1994 gathered results on the temperature-size rules phenotype in ectotherms from extensive number of studies and showed that 83% of the studies described that colder temperature resulted in significantly bigger body size. The same pattern of increased size at lower temperature was also observed in many insects and arthropods for body size and egg size (Azevedo et al., 2002; Azevedo et al., 1998; Czarnoleski et al., 2017; Ellers & Driessen, 2011; Fischer et al., 2006; Klok & Harrison, 2013; Steigenga et al., 2005). Although allelic variants and genes have been found that play an important role in body size plasticity (Bochdanovits et al., 2003; Ghosh et al., 2013; Lafuente et al., 2018; Li et al., 2006), the genetic architecture underlying this phenomenon is not fully uncovered yet.

Nematodes are not exceptional to this phenomenon. For instance, the nematode *Caenorhabditis elegans,* showed a 33% larger body size when grown at 10°C compared to nematodes grown at 25°C (Van Voorhies, 1996) and other temperatures (i.e 24°C) (Gutteling et al., 2007b; Kammenga et al., 2007). Part of this phenotypic variation in lower-temperature-dependent body size was caused by natural genetic variation in the calpain-like protease *tra-3* (Kammenga et al., 2007). Furthermore, the laboratory wide-used CB4856 strain are naturally smaller than N2 strain when grown at standard laboratory temperature which is associated with the variation in *npr-1* allele (Andersen et al., 2014).

Overall, *C. elegans* is an attractive organism for studying the genetics of plasticity to temperature. Its small genome, rapid life cycle (3.5 days at 20°C), genetic tractability, and a wealth of available experimental data have made this nematode a powerful platform to study the genetics underlying complex traits (Gaertner & Phillips, 2010; Snoek et al., 2020). Besides, *C. elegans* can be maintained completely homozygous, produce many offspring (200-300 offspring per self-fertilizing hermaphrodite), and can be outcrossed with rarely occurring males (Petersen et al., 2015; Sterken et al., 2015; Gaertner & Phillips, 2010). Furthermore, there are many temperature-related trait differences between two widely used divergent strains: N2 and CB4856. More specifically, studies reported that CB4856 and N2 differed in their response to temperatures in several life-history traits such as time to maturity, fertility, egg size, body size, lifespan, and also in gene expression regulation (Gutteling et al., 2007a; Gutteling et al., 2007b; Jovic et al., 2017; Kammenga et al., 2007; Li et al., 2006; Rodriguez et al., 2012; Viñuela et al., 2011). Despite these findings, we still do not have a full overview of the loci that affect plasticity at a larger range of different temperatures.

To further elucidate the genetic architecture of temperature affected body size plasticity in *C. elegans*, we selected 40 RILs derived from N2 and CB4856 parents (Li et al., 2006) to study the plasticity and genetic regulation of body-size traits (body-size and some internal organs size) under four temperatures and two developmental stages. First, we sought to investigate the effect of temperature and developmental stages to the reaction norms of the body-size traits, correlation between body size-traits within and between temperature-developmental stages, as well as investigating genetic parameters (heritability and transgressive segregation) of body size-traits and body-size plasticity. Subsequently, we investigated the genomic regions underlying these body-size traits across temperature-developmental stage combinations and plasticity traits under three temperature ranges. We found 18 QTL of body-size traits at certain temperature and developmental stages and five plasticity QTL. Many of the QTL for different traits colocalized at the same position within temperatures suggesting a pleiotropic effect or close linkage. Furthermore, some of the plasticity QTL also colocalized with body-size QTL in certain temperature, suggesting a possibility of co-regulatory loci underlying plasticity traits and traits itself. Moreover, the colocalizing QTL across temperatures indicating a possible temperature sensitive regulatory mechanism.

## Materials and methods

### Mapping population

The mapping population used in this study consisted of 40 RILs from a 200 RIL population derived from crossing of N2 and CB4856. These RILs are chosen because they resembled the smallest set of population with highest genetic diversity. The RILs were generated by (Li et al., 2006) and most were genotyped by sequencing, with a genetic map consisting of 729 informative (indicating a cross-over) Single Nucleotide Polymorphism (SNP) markers (Thompson et al., 2015). The strain names and genotypes can be found in Figure S1.

We confirmed that long-range linkage, between markers on different chromosomes, was not present in the population, by studying the pairwise correlation of the genetic markers in the used population (Figure S2).

### Cultivation and experimental procedures

*C. elegans* nematodes were reared following standard culturing practices (Brenner, 1974). RILs were kept at 20°C before experiments and three days before starting an experiment a starved population was transferred to a fresh NGM plate. An experiment was started by bleaching the egg-laying population, following standard protocols (Brenner, 1974). After bleaching, nematodes were placed on fresh NGM plate seeded with *E. coli* OP50. From that point onward, RILs were grown at four different temperatures: 16°C, 20°C, 24°C, or 26°C. At two time-points of developmental stages (L4 and adult) per temperature, microscope pictures (Leica DM IRB, AxioVision) were taken of three nematodes per line per temperature that were mounted on agar pads. The timepoints were chosen such that L4 and young adult nematodes were photographed, this was confirmed by mid-L4 vulva shape and adult vulva-shape and germline. The exact times are indicated in the sample data file (Table S1).

### Trait measurements and calculations

The number of RILs subjected to treatments per developmental stage was 40, except for treatment of temperature 24°C at L4 stage, where 39 RILs were used. Per life-stage and temperature we took measurements of 3 replicate individuals per RIL, and 6 replicate individuals of the parental lines N2 and CB4856 (from two independent populations). This resulted in 1056 pictures.

To quantify traits, the pictures were loaded into ImageJ (version 1.51f) and traits were manually measured. In total, nine body-size traits were measured: (i) body length, (ii) width at vulva, (iii) length of the pharynx, (iv) width of the pharynx, (v) length of the isthmus, (vi) length of the buccal cavity, (vii) length of the procorpus, (viii) surface postbulb, and (ix) surface nematode. To convert the measurement data from pixels to milimeters (mm), a figure of scale (in mm) was loaded to imageJ. Subsequently, the resolution of *C. elegans* picture and the scale picture were equalized. Next, in ImageJ we determined how many pixels were represented by 0.1 mm. This step was repeated 10 times and the average value was taken as standard conversion scale from pixels to mm. We also calculated body volume (assuming the nematodes body resembles a tube) as

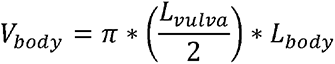

and the length/width ratio (L/W ratio) as the ratio of body length/width at vulva. For none of the traits we have a complete dataset due to difficulties in obtaining accurate measurements, the number of missing values for each trait are as follows: body length = 67; width at vulva = 102; length pharynx = 76; length isthmus = 220; surface postbulb = 219; surface nematode = 232; length buccal cavity =193; length procorpus = 242; body volume = 135; width pharynx = 65; length/width ratio = 135. All raw data can be found in Table S2.

### Analytical Software Used

Phenotypic data was analyzed in “R” version 3.5.2×64 using custom written scripts (R core Team 2017). The script is accessible via Gitlab: https://git.wur.nl/published_papers/maulana_2021_4temp. R package used for organizing data was the tidyverse (Wickham et al., 2019), while all plots were made using ggplot2 package (Wickham, 2011), except for heatmaps in Figure S3 which were made using the “heatmap ()” function provided in R. The data was deposited to WormQTL2 where it can be explored interactively (www.bioinformatics.nl/WormQTL2) (Snoek et al., 2020).

### Correlation analysis

The correlation between the traits in all treatment combinations was determined by the Pearson correlation index and plotted in a correlation plot. To correct for the effect of outliers (effect of very high or low value of single observation), we normalized the data as follows:

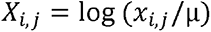

where *x* is individual observation of the traits in temperature *i* (16°C, 20°C, 24°C, 26°C)) and developmental stage *j* (L4, adult) while µ is the mean value of all traits.

### Transgressive segregation

To determine transgressive segregation of the traits among RILs panel, we performed multiple t-tests comparing all RIL panel to both parents for all traits per temperature and developmental stages. Transgression was defined when the traits of individual RIL is significantly different than both parents (p.adjust with FDR < 0.05; equal variance not assumed).

### Heritability estimation

Broad-sense and narrow-sense heritability of the phenotypic traits over RIL lines was calculated using Restricted Maximum Likelihood (REML) model to explain variation of the traits across the RIL lines (Kang et al., 2008; Rockman et al., 2010). The broad sense heritability was calculated according to the following equation:

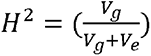

where *H^2^* is the broad-sense heritability, *V_g_* is the genotypic variation explained by the RILs, and *V_e_* is residual variation. The *V_g_* and *V_e_* were estimated by the lme4 model x_norm_ ∼ 1 + (1|strain) (Bates et al., 2015). The input data was normalized using log_2_ phenotype value to better fit the normality assumption.

Narrow-sense heritability is defined as the total variation in the population which is captured by additive effects. We calculated these using the heritability package in R, which estimates narrow-sense heritability based on an kinship matrix (Kruijer et al., 2014). The kinship matrix was calculated using the kinship function from the Emma package in R (Carta et al., 2011).

The significances of broad and narrow-sense heritability were determined by permutation analysis where the traits values were randomly assigned to the RILs. Over these permutated values, the variation captured by genotype and residuals were then calculated. This permutation was done 1,000 times for each trait. The result obtained were used as the by-chance-distribution and an FDR= 0.05 threshold was taken as the 50^th^ highest value.

### QTL mapping

QTL mapping was performed using custom script in R using fitted single marker model as follows:

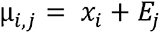

where µ is the averaged of all strains replicates in terms of their body size-traits *i*, of RIL *j* (N = 40) on marker location x (x= 1, 2, 3, …., 729).

Detection of QTL was done by calculating a -log_10_(p) score for each marker and each trait. To increase the detection power, all the values were log_2_ normalized mean trait values in temperature. To estimate empirical significance of -log_10_(p), the traits were randomly permutated value over the RILs 1,000 times. The calculation resulted in a significance threshold with false discovery rate (FDR) = 0.05 at a -log_10_(p) of 3.4 for QTL detection.

### Trait plasticity calculation

We divided plasticity ranges into three adjacent temperature groups: 16°C to 20°C, 20°C to 24°C, and 24°C to 26°C. Trait plasticity was defined as ratio between the trait mean value per nematode strain at 16°C to 20°C, 20°C to 24°C, and 24°C to 26°C.

### Heritability estimation of trait plasticity

Broad-sense and narrow-sense heritability of trait plasticity was calculated using the same model as in phenotypic traits heritability above. The broad sense heritability was calculated according to the following equation:

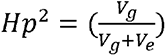

where *Hp^2^* is the broad-sense heritability of trait plasticity, *V_g_* is the genotypic variation explained by the RILs, and *V_e_* is residual variation. The *V_g_* and *V_e_* were estimated by the lme4 model x_norm_ ∼ 1 + (1|strain) (Bates et al., 2015).

The narrow-sense heritability was estimated based on kinship matrix calculated using the kinship function from the Emma package in R (Carta et al., 2011). The significances of broad and narrow-sense heritability were determined by permutation analysis as in phenotypic heritability estimation. The calculation was done for all temperature ranges (16°C to 20°C, 20°C to 24°C, and 24°C to 26°C) in adult and L4 stage.

### QTL mapping for trait plasticity

Plasticity QTL mapping was performed using fitted single marker model as follows

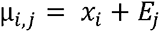

where µ is the averaged of all strains replicates in terms of their body size-traits *i*, of RIL *j* (N = 40) on marker location x (x= 1, 2, 3, …., 729).

Plasticity QTL detection was done by calculating a -log_10_(p) score for each marker and each trait. The calculation was done per temperature ranges. Empirical significance of - log_10_(p) was estimated by randomly permutated value over the RILs 1,000 times. The false discovery rate (FDR) = 0.05 at a -log_10_(p) of 3.0 was found as the significant threshold for plasticity QTL detection.

### Statistical power calculation

To determine the statistical power of our QTL and plasticity QTL dataset at the set threshold, we performed power analysis using the genetic map of the strains (n = 40) used per condition as in (Sterken et al., 2017). We simulated ten QTL per marker location that explained 20-80% of the variance, with increments of 5% (20%, 25%, 30%, …, 80%). Random variation was introduced based on a normal distribution with σ = 1 and µ= 0. Peaks were simulated according to effect-size, for example, a peak corresponding to 20% explained variation was simulated in this random variation. Based on the simulation, we analysed the number of correctly detected QTL, the number of false positives as well as undetected QTL. In addition, the precision of effect-size prediction and the QTL location were determined. The threshold used was based on 1000x permutation analysis -log_10_(p) > 3.4 for individual QTL and - log_10_(p) > 3.0 for plasticity QTL. The results of this calculation is presented in Table S5.

## Results

### *C. elegans* body size traits vary across temperatures and developmental stages

To investigate the impact of different genetic background, ambient temperature condition, and developmental stages on the body-size traits, we used a panel of 40 RILs derived from a cross between Bristol strain (N2) and Hawaiian strain (CB4856) (Figure S1 and S2) (Li et al., 2006). Each individual RIL was grown under four different temperature regimes (16°C, 20°C, 24°C, and 26°C). Once reaching the L4 and adult stage, we took pictures of each RIL with three individual replicates per strain and determined the body size parameters using ImageJ (Figure 1A; see materials and methods).

**Figure 1.**
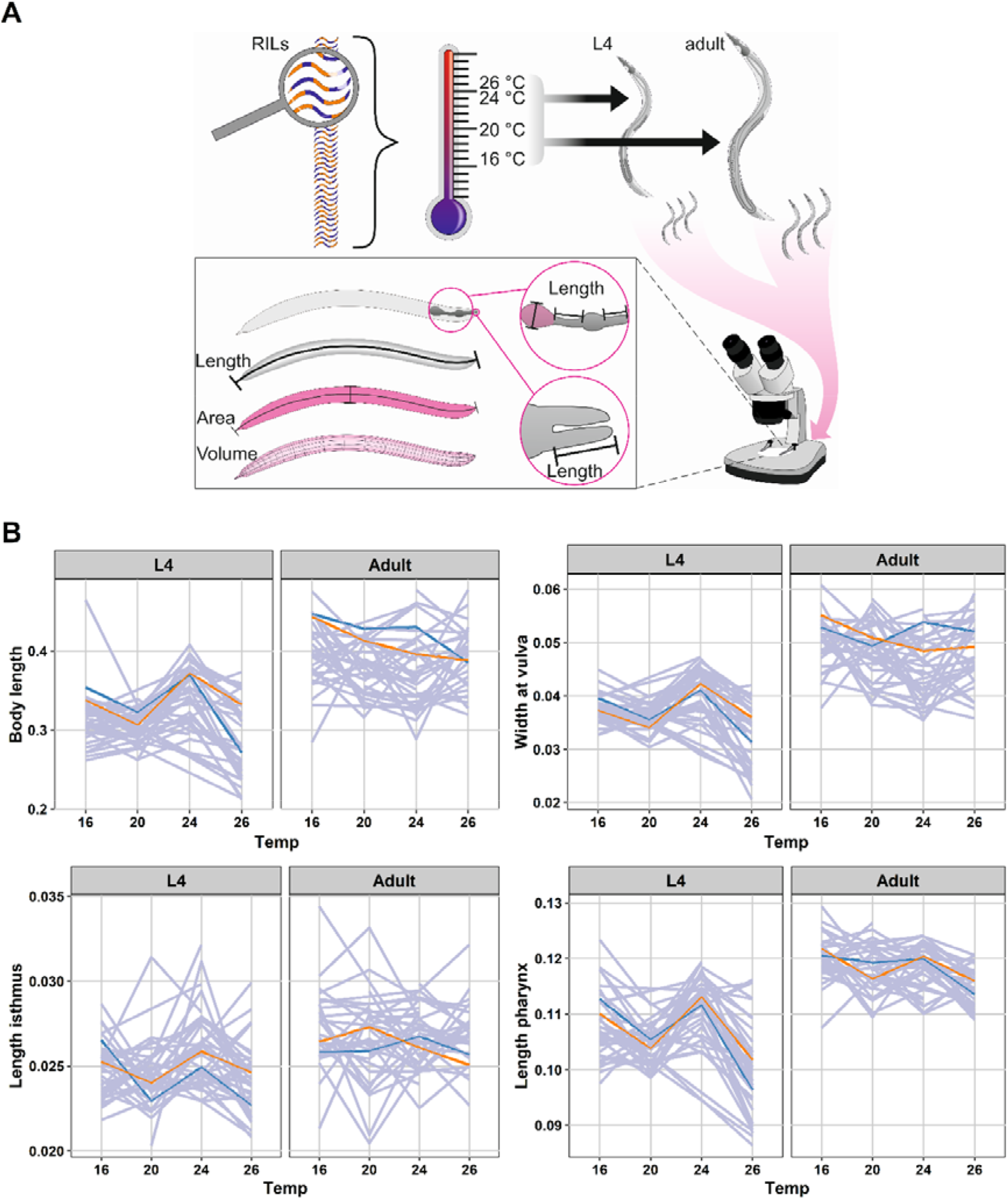
**(A) Flowchart of experimental overview**. A set of 40 RILs were grown at temperature 16°C, 20°C, 24°C, and 26°C. Separately, at L4 and adult stage, individual RIL with 3 replicates per RIL were photographed under microscope. Subsequently, the body-size traits of the RILs were determined using ImageJ. **(B) Reaction norms of four *C. elegans* body-size traits plasticity** across different temperature and developmental stages. The x-axis represents temperatures used while y-axis represent the mean value of the individual strains in their respective traits (in mm). Both parents are depicted in blue (CB4856) and orange (N2), while the RILs are grey.

The eleven body-size traits showed a dynamic variation across RILs when measured in different temperatures and developmental stages (Figure 1B; Figure S3), suggesting that these traits are indeed plastic. In general, we observed similar dynamic pattern of traits plasticity across RILs in L4 stage including N2 and CB4856. The trait values dropped from 16°C to 20°C, then increased from 20°C to 24°C, and dropped again from 24°C to 26°C. On the other hand, the trait values in adult stages show no similar pattern across the RILs, suggesting that the traits plasticity in adult stage are more sensitive to genetic background, whereas in the larvae the environment seems to play a larger role (Figure 1B; Figure S3). For several major body-size such as body length, body volume, width at vulva, and surface area of the nematodes of adult worms, we found that CB4856 did not completely follow the temperature-size rules (a decrease curve from 16-20°C, followed by an increase from 20-24°C, and decreased from 24-26°C) whereas Bristol N2 consistently grew bigger at lower temperatures. This gives a new insight to the finding from previous study (Gutteling, et al., 2007b; Kammenga et al., 2007), where CB4856 found to deviate the temperature-size rule, which that’s not always the case. This results shows that N2 body size was less plastic than CB4856.

To get insight into the relations between the traits measured, we performed a correlation analysis for all pairs of traits at the two developmental stages. We found that the level of between trait-correlation differed between L4 and adult stage, where temperature seems to be the main driving factor (Figure S4). Both in L4 and adult stage, the body-size traits displayed a strong positive correlation within the same temperature, and strong negative correlation between different temperatures, suggesting that the variation in the body-size traits were temperature specific. Interestingly, both in L4 and adult stage, the body-size traits of worm grown in 16°C and 26°C were separated into several small clusters, while the traits from 20°C and 24°C treatments formed a single positively correlated cluster. These results indicated that there were more similar patterns of variation over RILs in temperature 20°C and 24°C. Strongly correlated body-size traits imply that the same quantitative trait loci could be detected for these traits due to similar patterns of variation in the RILs, temperatures, and developmental stages.

To explore the source of variation of the body-size traits in the RILs population, we used principal component analysis (PCA) (Figure S5). The PCAs describes the variation of the traits based on temperatures and genetic background per developmental stage. At the L4 stage, the first principal component captured 45.5% of the variation where the 16°C temperature animals were most distal from the other temperatures, while the second principal component captured 24% of the variation where the 24°C and 26°C temperatures were most distal. We found that at L4 stage, the RILs were more similar in lower temperature (16°C) while in 20°C, they were distributed across the PC plot. Subsequently, the value of body-size traits of the nematodes at 24°C were similar to the values at 26°C, but divided into two cluster (Figure S5). On the other hand, the individual RILs did not show any clear clusters at adult stage, indicating there was high variation between the RILs as a result of interaction between environment and the genetic background. This result combined with the correlation analysis show that there was a substantial variation in the RILs, suggesting that it was possible to detect QTL controlling the traits.

### Transgressive segregation and heritability indicate a complex genetic architecture underlying body-size traits

Upon inspecting the distribution of trait variation in the RILs compared to N2 and CB4856, we observed high levels of variation exceeding those of the parental strains (Figure S6). This suggests transgressive segregation within the RIL population. Hence, we tested the trait values of each RIL versus the parents. We found transgression for almost all traits per temperature-developmental stage combinations (t-test, p.adjust FDR < 0.05) (Table S3). Our findings show that the number of two-sided transgressive RILs depended on the combination of temperature and developmental stage (Figure 2A Figure 2B; Table S3). Whereas in the L4 stage the number of transgressive RILs was constant under 16°C, and 20°C, slightly dropped under 24°C – and then increased at 26°C. Conversely, in the adult stage, the number of transgressive strains decreased as the temperature increased. Moreover, it shows that the parental lines have both positive and negative alleles that interact with the environment leading to a more robust/stable phenotype over a broader temperature range. Using ANOVA, we found that developmental stage was indeed the factor driving transgression (p = 0.0275; R^2^ = 0.073 Table 1) whereas temperature alone showed no relation to the transgression (p = 0.786; R^2^ = 0.015). These findings indicate environment and age-specific effects on the regulation of body-size traits.

**Figure 2.**
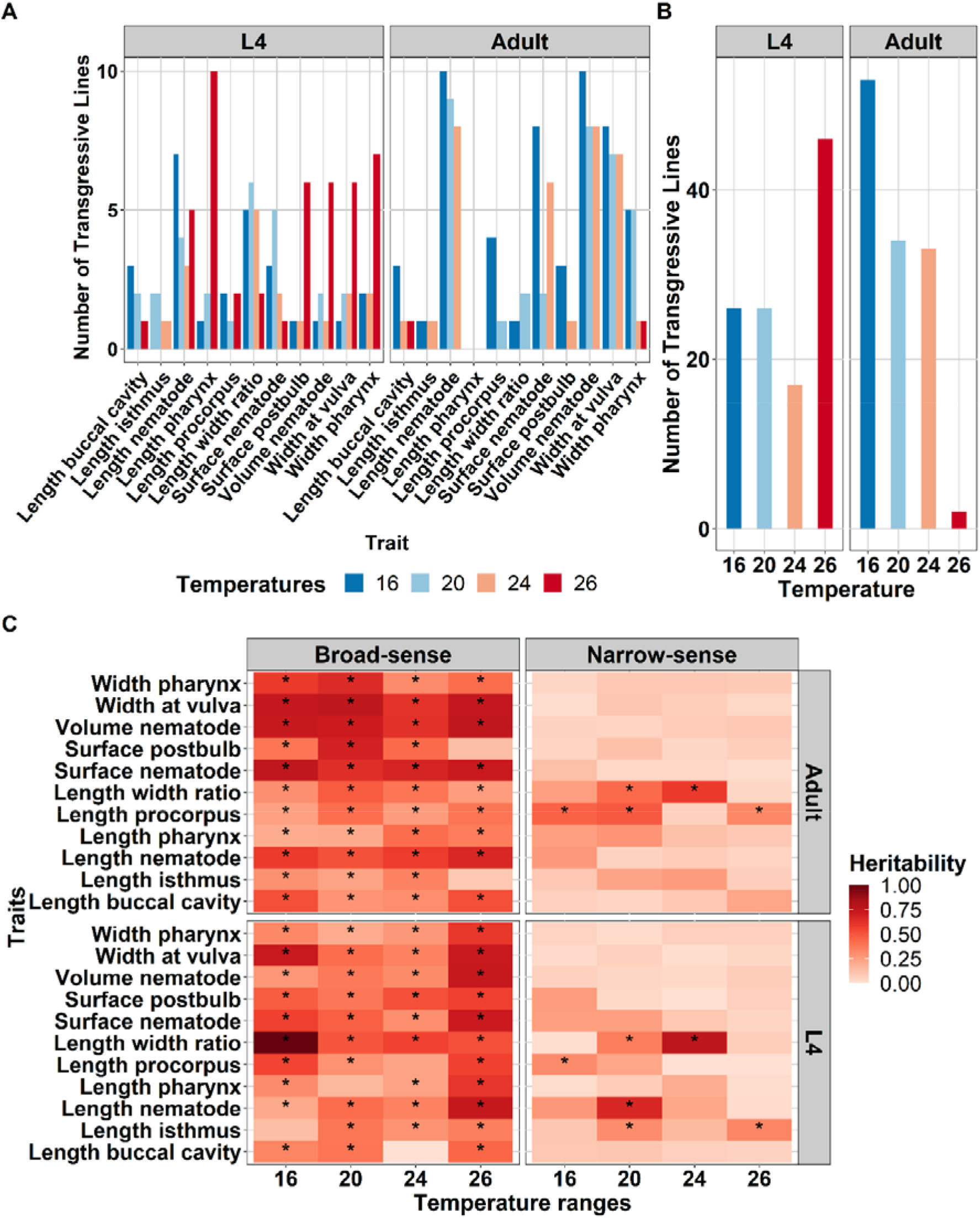
**(A) The number of transgressive lines of eleven body-size traits** in *C. elegans* across temperatures and developmental stages. The traits are on the x-axis while y-axis represents the number of transgressive lines based on multiple t-test of traits in each individual lines against both parents (p.adjust FDR < 0.05). Colours represent the temperatures treatment, corresponding to the legend on the bottom side. **(B) The number of transgressive traits found in the RIL population** per treatments combination (temperature-developmental stage). The temperature is on the x-axis while y-axis represents how many transgression found within those temperatures. Developmental stages are depicted on the above side of the graph. **(C) Broad-sense and narrow-sense heritability** of body-size traits across temperature and developmental stages. On the x-axis are temperature and on the y-axis are the traits measured. The colour gradient represents the heritability values as depicted on the legend on the right side of the plot. Asterisk (*) inside the box indicate significant heritability values (FDR0.05 based on 1000 permutations). Developmental stages are depicted on the right side of the plot whereas the types of heritability are on the top.

**Table 1.**
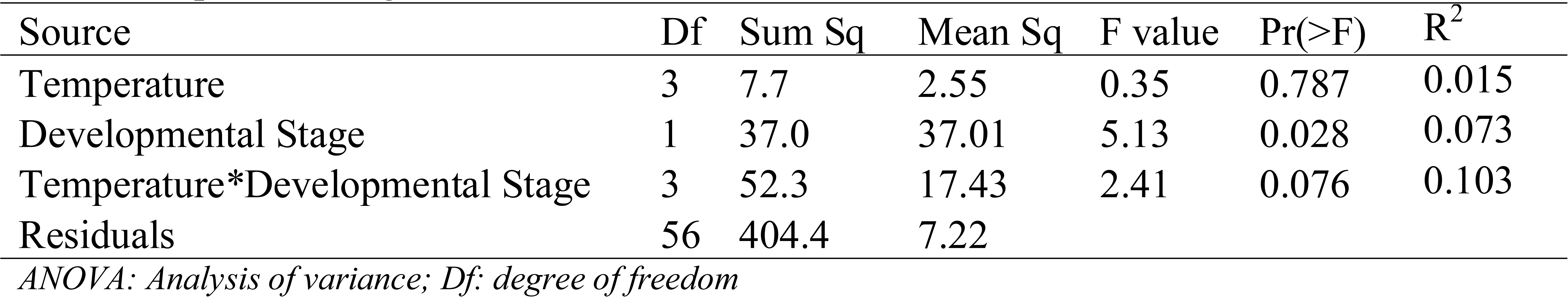
Results of ANOVA for the number of transgressive lines over temperatures and developmental stages.

Next, to determine the proportion of variance in body-size traits that were caused by genetic factors, we calculated the broad sense-heritability (*H^2^*) of each trait. We found significant heritability (REML, FDR = 0.05) for 81 out of 88 traits in developmental stage-temperature combinations. The significant heritability ranged from 0.0 (length buccal cavity at 24°C in L4) to 0.99 (length/width ratio at 16°C in L4) (Figure 2C; Table S4). Hence, for a large fraction of traits we could detect a high contribution of genetic factors. In addition to *H^2^*, we calculated the narrow-sense heritability (*h^2^*) to identify how much of the variation could be explained by additive allelic effects. This analysis suggested that there were 11 traits with significant additive effect (REML, FDR < 0.05; Table S4). For nearly all body-size traits we detected *H^2^* well beyond *h^2^*, indicating a role for epistasis in the genetic architecture of the traits.

To understand the contribution of temperature and developmental stage on heritability of all traits measured, we conducted an ANOVA (Table 2). The results suggested a trend that the main factor driving *H^2^* was temperature (*R^2^* = 0.008, p = 0.078) and its combination with developmental stage (*R^2^* = 0.084, p = 0.066). On the other hand, developmental stage showed little relation to the variation of *H^2^ (R^2^ =* 0.005, p = 0.493). In the adult stage, we observed there were four traits (width at vulva, body length, body volume, and surface area of nematodes) which *H^2^* are relatively robust across all temperatures while in L4 they were found to be more variable across temperature. These four traits, were affecting each other and observed to be positively correlated (Figure S3). Taken together, overall body-size traits show significant *H^2^*, indicating a substantial effect of the genetic background on the variation in these traits in this population. Moreover, the correlation between some traits indicate a shared genetic architecture between the traits These results indicate a higher chance of detecting QTL on the traits measured.

**Table 2.**
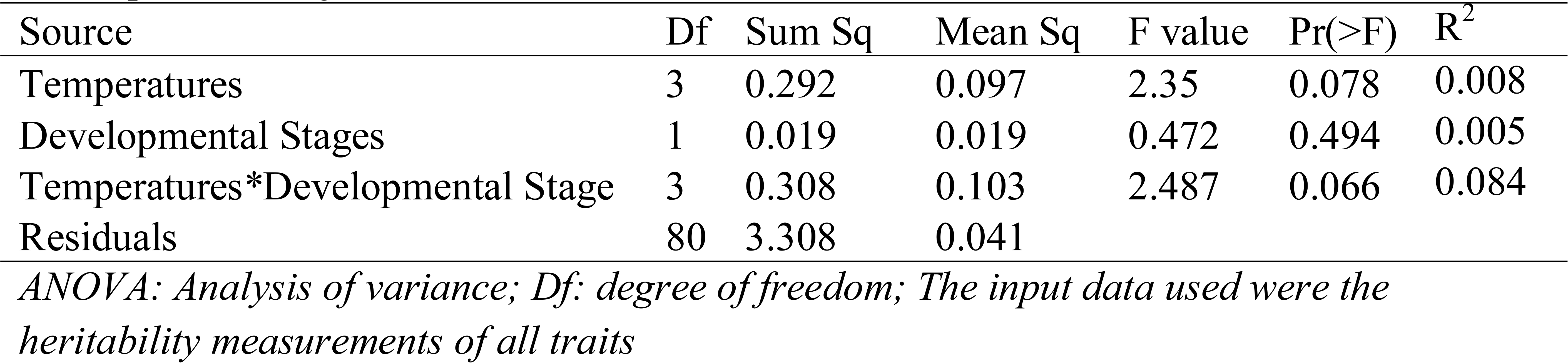
Results of ANOVA for *H^2^* of all traits over temperatures and developmental stages

**Table 3.**
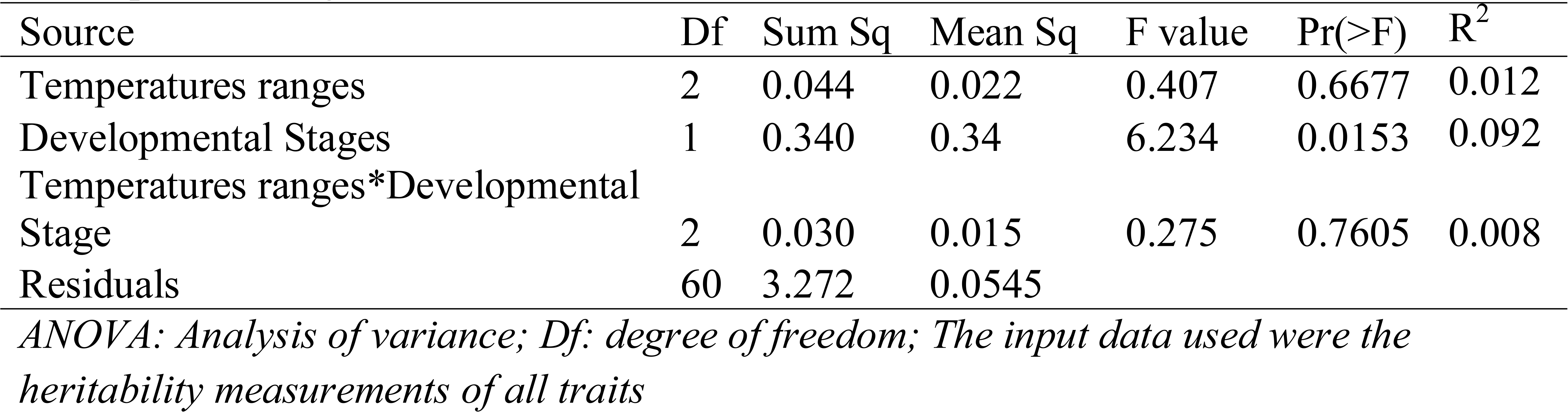
Results of ANOVA for *H^2^* of all trait plasticity over temperatures and developmental stages

### QTL underlying body-size traits in *C. elegans* are influenced by temperature and developmental stages

To identify underlying loci controlling the variation of body-size traits, we performed QTL mapping for all the body-size traits measured in the 40 RILs. Analysis of statistical power showed that our population can detect 80% of true QTL explaining 60% of the variance (Table S5). Using log-normalized mean values per RIL as input, we found 18 significant QTL (-log_10_(p) = 3.4, FDR = 0.05) with -log_10_(p) scores ranging up to 6.5 in each temperature and developmental stages (Figure 3, Table S6). We found QTL explaining 28-53% of variance among the RILs (Table S6). We found 7 QTL in the L4 stage namely surface area, length pharynx, body length, length procorpus (detected at 20°C), length/width ratio (detected at 20°C and 24°C), and surface postbulb (detected at 16°C) . For the adult stage, 11 QTL were detected for the body-size traits. Here, we found QTL evenly distributed over the temperatures: two at 16°C, two at 20°C, three at 24°C, and four at 26°C (Figure 3A). Of the 18 significant QTL, eight were located on chromosome X, five QTL on chromosome V, three on chromosome I and two on chromosome IV.

**Figure 3.**
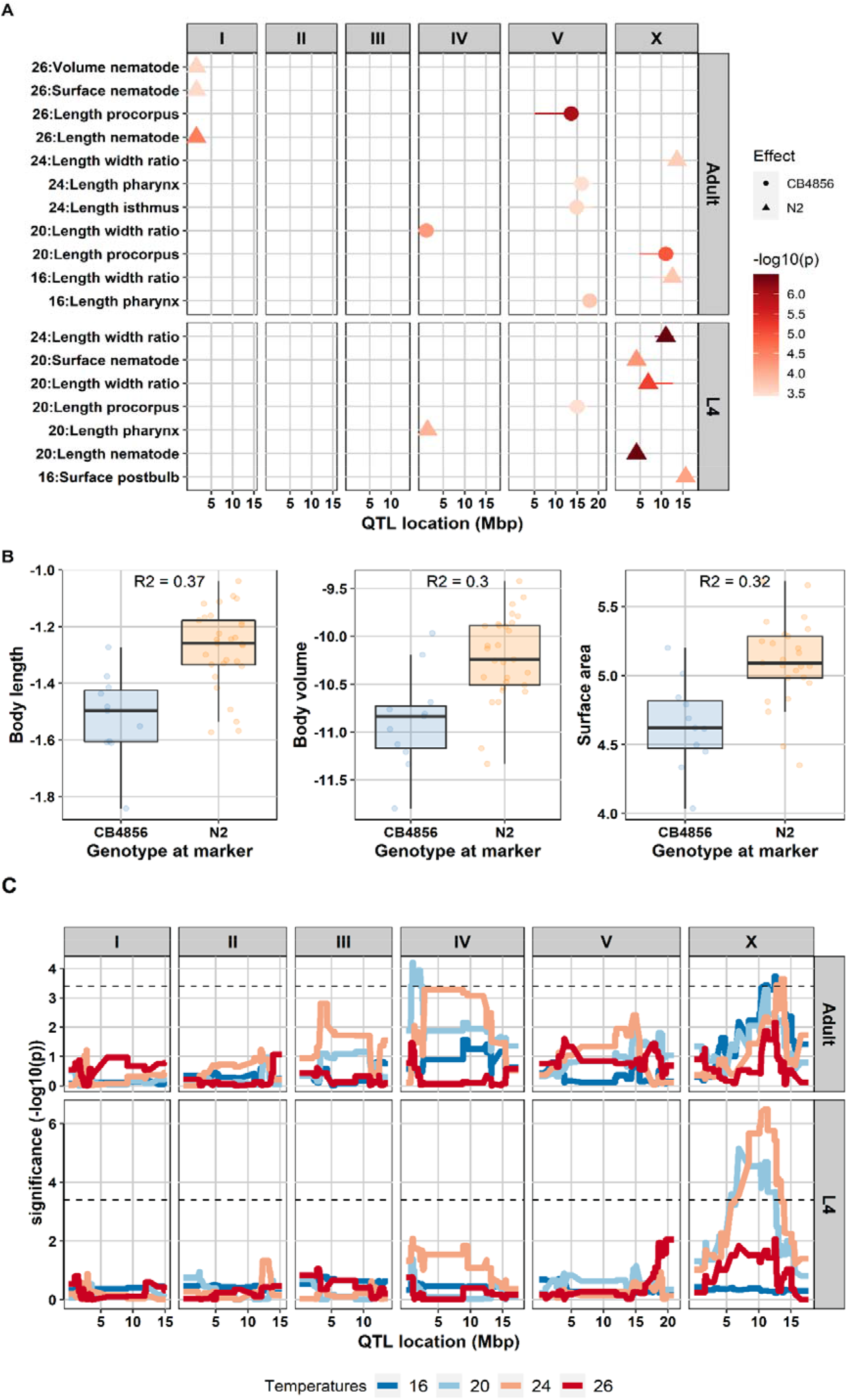
**(A) QTLs found for body-size traits in the 40 RILs.** The x-axis represents the position of the QTL in mega base pairs (Mbp) for each chromosome and y-axis displays the corresponding significant QTL from a single marker model. In total 18 QTL were found with -log_10_(p) score ranging from 3.44 to 6.49 (-log_10_(p) threshold 3.4, FDR = 0.05). Shapes represent genotype effect: dots = CB4856; triangles = N2. **(B) Allelic effects of QTL** for body length, body volume, and surface area of the nematode in adult stage at 26°C at the peak marker location. RILs that have N2 marker at this locus relatively have bigger body size compared to those that have CB4856 marker. The genetic variation on chromosome I can explain 30 to 37% of the variation in body length, body volume, and surface nematode at that condition. **(C) QTL profile of length/width ratio**. The QTL analysis were performed across four temperatures (16°C, 20°C, 24°C, 26°C) and two developmental stages (L4 and adult) using a single marker model. X-axis displays genomic position in the chromosome corresponding to the box above the line while y-axis represents the - log_10_(p) score. Box in the right graph show the developmental stages. Blue line represents QTL at 16°C, light blue line at 20°C, orange line at 24°C, and red line at 26°C. Black-dash line represents -log_10_(p) threshold (FDR = 0.05).

We observed QTL-hotspots for various traits. For example, the chromosome I QTL (surface area, body volume, and body length) were positively correlated traits, mapped in the same developmental stage and temperature combination. Hence, this could point to a body-size QTL, where the N2 genotype was associated with larger body size compared to the CB4856 genotype (Figure 3B). Interestingly, all QTL on chromosome V were associated with the size of the feeding-apparatus, were found over various temperatures, and were all associated with an increased size in CB4856 (Figure 3A; Figure S7). In contrast, traits related to the overall body size (e.g. volume) were almost exclusively associated with an increased size due to the N2 allele (Figure 3B).

In line with the indications of the correlation- and heritability analyses, we found evidence for environment (temperatures), age (developmental stage) and genotype interactions. For example, for length/width ratio (Figure. 3C) in the adult stage, significant QTL in chromosome X were detected for the worms grown at 16°C and 24°C, one QTL on chromosome IV for worms grown at 20°C, and no significant QTL detected at 26°C. When we mapped the trait in adult worms grown at 16°C, we found a significant QTL on chromosome X which we did not find in the L4 stage at 16°C. The same result was found for QTL at temperature 20°C at adult stage on chromosome IV which was not present in L4 stage. Similar patterns of (dis-) appearance were observed for many traits (Figure S7). Hence, traits may be regulated by different set of genes (loci) dependent on temperature-environment and developmental stage. This indicates that there is a considerable effect of genotype-environment interactions.

### The RIL population revealed plasticity QTL for several body-size traits

Phenotypic plasticity is the change of the expressed phenotype in different environments. To determine the amount of variation in body-size plasticity was due to genetic factors, we calculated the *H^2^* of each set of neighbouring experimental temperatures: we defined plasticity as the ratio of the traits in 16°C to 20°C, 20°C to 24°C, and 24°C to 26°C. We found significant *H^2^* (REML, FDR = 0.05) for trait plasticity for 45 out of 66 traits in developmental stage-temperature ranges combinations. In contrast, for *h^2^* of traits plasticity, there were only three traits (length pharynx, length procorpus, and width pharynx, ad adult stage on temperature ranges of 16°C to 20°C) with significant additive effect (REML, FDR < 0.05; Table S7). The significant *h^2^* values ranged from 0.19 (length pharynx at adult stage on 20°C to 24°C range) to 1.00 (length/width ratio on 16°C to 20°C in L4 stage) (Figure 4A; Table S7). Consistent with the patterns observed across temperatures in the L4 versus the adult stages (Figure 1B), there were fewer significant broad- and narrow-sense heritabilities observed in L4 stage compared to the adult stage, indicating higher environmental variation of traits plasticity in L4. In summary, we observed a strong effect of developmental stages to the heritability of trait plasticity which was also strongly dependent on the body-size trait under study.

**Figure 4.**
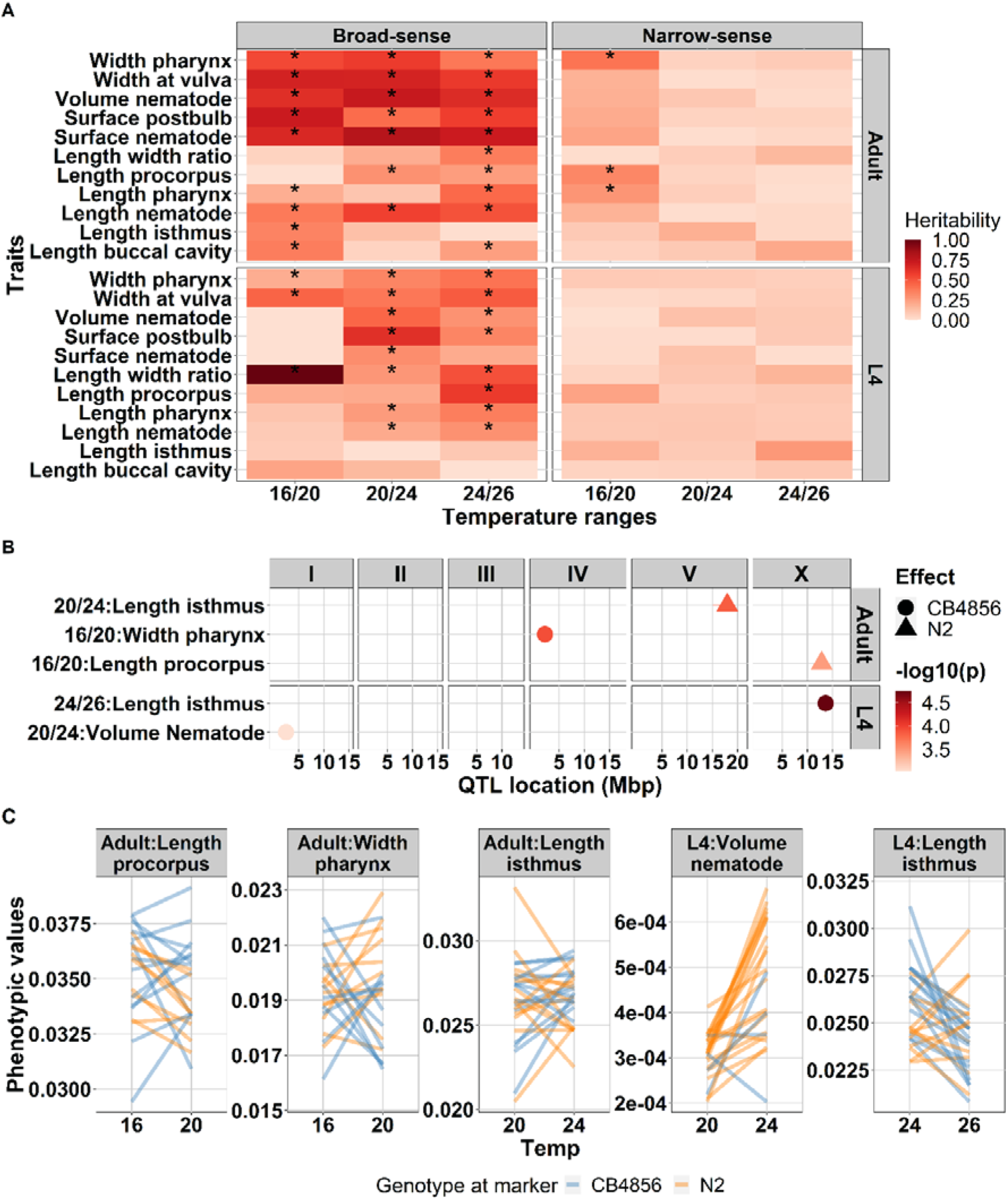
**(A) Broad-sense and narrow-sense heritability** of body-size traits plasticity across temperature range and developmental stages. On the x-axis temperature plasticity and y-axis are the traits measured. Colour gradient represent the heritability value as depicted on the legend on the right side of the plot. Asterisk (*) inside the box represent significant heritability value (FDR0.05 based on 1000x permutation). Developmental stages are depicted on the right side of the plot whereas the type of heritability are on the top. **(B) Plasticity QTLs found for body-size traits in the 40 RILs.** The x-axis represents the position of the QTL in million base pairs (Mbp) for each chromosome and y-axis displays the corresponding significant plasticity QTL based on a single marker model. In total five plasticity QTL were found with -log_10_(p) score ranging from 3.03 to 4.75 (-log_10_(p) threshold 3.0. **(C) Phenotypic values (in mm) of the corresponding plasticity QTL.** X-axis represents the temperature regime where the plasticity QTL identified. Y-axis represents the mean phenotypic values of the traits. Orange lines were RILs with N2 genotype at the peak marker location, whereas blue lines represent RILs with CB4856 genotype at the marker location. The traits are depicted on the top of each plots.

In contrast to *H^2^* values per temperature-trait combination where the main factor driving the heritability was temperature, in *H^2^* of plasticity, developmental stage (*R^2^ =* 0.092, p = 0.0153) was the main explanatory factor. On the other hand, temperature ranges showed little relation to the variation of plasticity *H^2^ (R^2^ =* 0.012, p = 0.667). Taken together, overall body-size traits plasticity showed significant *H^2^*, indicating a substantial effect of the genetic background on the variation of these traits in this population.

The previous results suggested that QTL affecting body-size traits can be located on different chromosomes when measured in different environment, indicating an environment-QTL interaction. To further understand the mechanism of trait plasticity, we mapped QTL for trait plasticity. Statistical power analysis indicated that we can detect 80% of QTL explaining 60% of variation (Table S5). For all three conditions, we found five significant plasticity QTL. Two plasticity QTL were found in the temperature range from 16°C to 20°C harbouring a locus associated with width pharynx at adult stage, and length of the procorpus at adult stage. Two plasticity QTL in temperature range of 20°C to 24°C were associated with length isthmus at adult stage and body volume at L4 stage. One plasticity QTL in temperature ranges of 24°C to 26°C related to length isthmus at L4 stage (Figure 4B, 4C; Table S8). Hence, we found less QTL than for the individual temperatures. However, this was to be expected since the narrow-sense heritability of plasticity was lower and resulted in fewer significant values.

We compared the plasticity QTL to the QTL mapped for the individual temperatures. Of the five plasticity QTL, three of them were colocalized with QTL for body-size traits within individual temperatures (Figure 3A; 4B): (1) plasticity QTL of length procorpus in the temperature range from 16°C to 20°C colocalized with QTL for length width ratio of adult nematode at 16°C; (2) plasticity QTL for length isthmus of L4 worms in temperature ranges of 24°C to 26°C colocalized with length width ratio of adult worms in 24°C; (3) plasticity QTL for length isthmus of adult worms in the temperature range of 20°C to 24°C colocalized with QTL for length pharynx at adult stage at 16°C. These results indicate that although plasticity can be reflected in individual temperatures, contrasting trait values over varying conditions reveals new insights into the underlying loci.

We continued by investigating the direction of the plasticity QTL. In two of the QTL the N2 genotype had a negative effect on the trait value, namely for length procorpus (16°C to 20°C at adult stage) and plasticity QTL for length isthmus (20°C to 24°C at adult stage). In other words, the RILs harbouring N2 genotype at the peak marker location have decreased phenotype, while CB4856 genotype have an increased phenotype (Figure 4C). In contrast, for plasticity QTL of width pharynx (16°C to 20°C at adult stage), body volume (20°C to 24°C at L4 stage), and length isthmus (24°C to 26°C at L4 stage), the RILs with an N2 locus display an increase of the trait value. The slope of reaction norm of the trait plasticity indicates allele(s) that affect the trait variation both in environment 1 and environment 2 linked to genetic variation in plasticity (Lafuente & Beldade, 2019). Together, our results indicate that temperature-related phenotypic plasticity of body-size traits were not governed by alleles with large changes in effect-sizes over the temperature gradient as we map few QTL. The heritability analysis indicate that in general, temperature-related plasticity is regulated by complex genetic effects over the course of the gradient.

## Discussion

### Body-size traits reaction norm reveals a genotype x environment interaction

In most ectotherms, temperature is an important factor driving body size and is related to life history traits (Angilletta & Dunham, 2003; Ellers & Driessen, 2011; Ghosh et al., 2013; Peng et al., 2007). This was also found for *C. elegans* (Gutteling et al., 2007a,b; Kammenga et al., 2007). In this study, we used a *C. elegans* RIL population to study the underlying genetic regions that regulate the body-size traits, both main body traits (i.e length, width at vulva, volume, length/width ratio, and surface area) as well as internal organs and feeding apparatus (i.e length isthmus, length procorpus, length pharynx, width pharynx, and surface postbulb), and the plasticity of such traits in different temperatures at two developmental stages.

We observed that the reaction norm of the body-size traits over temperatures were varied depending on the trait and individual genotypes and showed a clear genotype-environment interaction (GxE) (Beldade et al., 2011; Lafuente & Beldade, 2019; Lafuente et al., 2018; Saltz et al., 2018). The fact that some RILs do not follow the temperature-size rule, especially for major body-size traits, might be the results of CB4856 genotype in the RILs, as this strain is known to deviate from this rule (Gutteling et al., 2007; Kammenga et al., 2007). Interestingly, we also found that the type of reaction norm was affected by developmental stage of the animal. For most of the traits measured, adult worms displayed non-parallel reaction norm across temperature, as opposed to L4 stage worms which showed relatively similar non-linear parallel reaction among genotypes in most of the traits (Figure S4). This indicates that variation in phenotypic plasticity was not common in juvenile animals suggesting the absence of (or less significant) GxE for most genotypes as described in (Saltz et al., 2018). This differences of L4 and adult stage reaction norm could stem from differences in interaction between gene-expressions and environment (temperature) in L4 and adult (Lafuente & Beldade, 2019; Li et al., 2006; Snoek et al., 2017; Viñuela et al., 2010).

Within *C. elegans* it was previously known that the reaction norm of body length and body volume was defied in the CB4856 strain due to a polymorphism in *tra-3* (Gutteling et al., 2007b; Kammenga et al., 2007). This was measured over a two-temperature gradient from 12-24°C. We found that this not always the case, as the body size of CB4856 was dynamics (decrease and increase) over the course of temperatures. Meanwhile, for N2, we found that the negative linear norm was most apparent from 16 to 20, and at higher temperatures the overall body-size (e.g. length, width at vulva, surface area, and volume) were robust. It suggests that the body-size trait of N2 follows the “threshold” reaction norm model, instead of the linear function as described in (Lafuente & Beldade, 2019). This discrepancy could only be revealed by using four temperatures (16°C, 20°C, 24°C, and 26°C) as in this study.

Previous studies have suggested the effect of temperatures on genetic correlation of body size and several life history traits in different species. Lafuente *et al*., (2018) found that body size (thorax and abdomen) of *D. melanogaster* reared at 17°C and 28°C significantly correlates with chill coma recovery and survival of *Metarhizium anisopliae* fungi. Norry & Loeschcke (2002) found a positive correlation effect of lifespan with temperature and sex in *D. melanogaster* at 25°C where the male flies lived longer. However, this effect was reversed at 14°C. In *C. elegans*, an 18% of increase lifespan due to heat-shock was reported for CB4856 but not for N2, whereas the RILs showed a wide range of lifespan variation (Rodriguez et al., 2012). Our results are in agreement of other previous studies (reviewed in Sgro & Hoffmann, 2004) that different environmental conditions result in a different correlation power, suggesting that evolutionary trajectories on trade-offs between traits, especially the traits that are controlled by specific loci, depend strongly on the environmental condition.

### Genetic parameters and QTL analysis indicate a complex genetic regulation of body-size traits

In a population derived from two diverse parents, it is common to detect extreme phenotypes exceeding way beyond the parents (transgression) (Rieseberg et al., 1999). Transgression can represent genetic complexity of a trait, for example due to genetic interaction (epistasis) or it could mean that the trait is controlled by multiple loci with opposite effect combinations in the parental strains resulting in a similar phenotype. Transgression has been reported for *C. elegans* life-history traits such as egg size, number of eggs, body length (Andersen et al., 2015; Gutteling et al., 2007b; Kammenga et al., 2007), lifespan (Rodriguez et al., 2012), as well as metabolite levels (Gao et al., 2018), and gene expression (Li et al., 2006). We found transgressive segregation for almost all traits in temperature and developmental stage combinations, indicating a complementary actions of multiple loci underlying these traits.

We then calculated the broad-sense heritability to investigate the proportion of variance explained by genetic factors in our RIL population. It should be noted that because of the necessity of using batches *H^2^* represents an upper-bound. Still, our estimation of broad-sense heritability of adult body length at 20°C (0.51) was similar (*H^2^* = 0.57) as reported by Andersen et al., (2014). In addition, the *H^2^* of body volume (0.71) and width at vulva (0.76) in this study are also similar to (Snoek et al., 2019), which were 0.77 and 0.75, respectively. Heritability is a population trait characteristic and highly depends on the type of population used and environment. Therefore, the fact that we found similar heritability with previous works indicate that the variation of these traits is quite stable between different mapping population. This could also mean that the relative effect of the micro-environment as well as the stochasticity is small. Furthermore, similar patterns of heritability that changed over temperatures (12°C and 24°C) were reported for body mass (volume), growth rate (change in body length), age at maturity, egg size, and egg numbers (Gutteling et al., 2007a; Gutteling et al., 2007b).

By QTL mapping, we found 18 significant QTL for 88 temperature and developmental stage combinations regulating body-size traits. Here, we showed QTL of some the traits were colocalized in the same location in chromosome. For example, body length and surface area of the nematode in L4 stage at 20°C shared the same genomic region on the left arm of chromosome X. This is expected since these traits have strong positive correlation. All the colocalized traits showed the same QTL effect where N2-derived loci were associated with an increase in size. It is possible that such co-localized QTL were the result of a single pleiotropic modifier affecting various aspects of the *C. elegans* physiology. On the other hand, this might be the result of unresolved separate QTL (Dupuis & Siegmund, 1999; Gutteling et al., 2007b; Sterken et al., 2020).

As body length at 20°C has been investigated across multiple studies, we used it to cross-reference our mapping. The same location (chromosome X: 4.9 Mb) was mapped in two other studies (Andersen et al., 2014; Andersen et al., 2015). Furthermore, many QTL located in left arm of chromosome X were associated with body length, indicating the alleles controlling these traits might the same or linked with alleles of body length. In other study, using a multi-parent RIL, it was found that loci located in the same position (chromosome X around 4.5 to 5Mb) was associated with length/width ratio which is also related to body length (Snoek et al., 2019). For the same trait, (Snoek et al., 2014) found the QTL in different chromosome (i.e chr IV), meaning that our study has the power to reveal the previous undetected QTL.

Nagashima et al., (2017) summarized factors and its genetic basis involved in regulating body size in *C. elegans* including *DBL-1, TGF-*β signalling, *DAF-2, rict-1, sma-5, wts-1, IGF*-signalling*, tra-3, npr-1, cat-2, dop-3, eat-2, pha-2,* and *pha-3*. We manually checked the position of those genes in www.wormbase.org and found that none of the genes located under our QTL, except for *npr-1* (wormbase, 2021). Our QTL in chromosome X overlapped with the location of the Neuropeptide receptor 1 (*npr-1)* allele, which encodes the mammalian neuropeptide Y receptor homolog. This allele is a known pleiotropic regulator affecting traits such as lifetime fecundity, body size, and resistance to pathogens mediated by altered exposure to bacterial food (Andersen et al., 2014; Nakad et al., 2016; Reddy et al., 2009; Sterken et al., 2015). Moreover, 7 out of 8 QTL that colocalized in chromosome X has an increase size that are associated with N2 genotype, which support our hypothesis that those seven trait could be *npr-1* regulated.

Although not significant, we found potential QTL of body volume and width at vulva of adult nematode at 20°C and 24°C on the left arm of chromosome IV (Figure S7) which overlapped with QTL identified previously for body volume by (Gutteling et al., 2007b; Kammenga et al., 2007) at 24°C using a larger population of RILs, also in chromosome X at 20°C using multi-parental RIL (Snoek et al., 2019). These results indicate that these QTL represent robust and predictable genetic associations with temperature and size.

From 18 significant QTL, 9/18 (50%) were transgressive and 15/18 (83%) of the QTL had moderate to high heritability (> 0.3). These findings indicate a highly complex genetic regulation of many body-size traits that could involve multiple interaction of different genetic variants. This was supported by the higher value of broad-sense heritability compared to narrow-sense heritability which suggests that the driving factors of most heritable traits were additive loci of opposing effects or genetic interactions.

### Mapping of plasticity increments indicates small effect-size changes resulting in shifting loci

By mapping phenotypic plasticity over adjacent temperatures, we only found five plasticity QTL. We found two plasticity QTL over 16°C to 20°C that were related to width pharynx and length procorpus, both in adult stage. In addition, we detected two plasticity QTL over 20°C to 24°C that was related to length isthmus in adult stage and body volume at L4 stage. Lastly, one plasticity QTL over 24°C to 26°C associated with length isthmus at L4 stage. These result suggests that the QTL associated plasticity was environment specific, meaning that the candidate genes in the QTL region are differentially expressed depending on environmental conditions (Gutteling et al., 2007b).

We found little overlap between QTL for trait plasticity and QTL of traits in specific temperatures and developmental stages. Moreover, the plasticity QTL and traits QTL that colocalized were related to different traits. This low overlap of plasticity QTL and body-size trait QTL was also reported for *D. melanogaster* (Lafuente et al., 2018). Our results contribute to the long standing debate on the genetic basis of plasticity (whether it is controlled via specific loci for trait plasticity or via the same loci that regulate trait at certain environment) (Via et al., 1995). We showed that genetic basis of trait plasticity, to some extent, can be the same with the genetic basis of traits in certain environment, which support both ideas in congruence with previous papers, e.g. (Têtard-jones et al., 2011). We also showed that one loci can be responsible for different traits as well as responsible for plasticity in different environments. These findings may also indicate an allelic sensitivity model underlying plasticity mechanism where loci display environmental-based allelic sensitivity (Scheiner, 1993). The fact that these plasticity QTL were colocalized with QTL of traits at certain environment may suggest that the QTL contains loci/alleles that are activated when the population in a different environment or in an unusual condition (Paaby & Rockman, 2014) and can points to the co-evolution of traits plasticity and traits at given environment.

## Acknowledgements

The authors thank Simone Ariens for measurements on the microscopy images and Miriam Rodriguez for technical assistance. We thank Anne Morbach (Schlaugemacht.net) for making Figure 1A. We thank Harm Nijveen for assistance with WormQTL2. We thank Lisa van Sluijs for the critical review of the draft manuscript. We thank Fred van Eeuwijk for helpful discussions.

## Availability of data and materials

The strains used in this study can be requested from the authors. The underlying data is included in the paper and interactively accessible via WormQTL2.

## Author contributions

MGS, LBS and JEK conceived and designed the experiments. MGS and JAGR conducted the experiments. MIM analyzed the data with input from MGS. MIM and MGS wrote the manuscript, with input from JEK and LBS. All authors commented on the manuscript.

## Supplementary material

**Supplementary Table S1**. An overview of the experimental set up and raw data (in pixels) of this study. Per nematodes individual, detailed workflow of the experiment starting from the nematodes line used, replicate per lines, bleaching date and time per nematodes, picture date and time per nematodes, developmental stage and age of the nematodes when the pictures were taken, and raw data of the traits measured (in pixel) are given.

**Supplementary Table S2**. Calculated raw data (in mm) and the mean value of every traits measured. Conversion of the raw data was done by calculating the conversion constant from pixels to milimiters using ImageJ. Mean values of the trait were calculated per nematodes lines per temperature and developmental stages. All blank cells containing no value were removed.

**Supplementary Table S3**. Calculated transgressive segregation. Per Strain, stage, temperature, and trait the t-test significance - adjusted for multiple testing by the false discovery rate (FDR) method – is given. When both the N2 and CB4856 parent were significant, a strain was considered to show transgression.

**Supplementary Table S4**. Calculated broad sense and narrow sense heritability. Per trait, developmental stage and temperature the broad-sense heritability *H^2^* was calculated (H2_REML), the value for the 0.05 FDR threshold is also given (H2_FDR) as is whether the *H^2^* was significant (H2_significance). Similar for the narrow-sense heritability *h^2^* was calculated (h2_REML), the value for the 0.05 FDR threshold is also given (h2_FDR) as is whether the *h^2^* was significant (h2_significance).

**Supplementary Table S5.** Analysis of power by simulation of QTL. Ten simulated QTL per marker location was performed that explained 20-80% of the variation, with an increment of 5% (20%, 25%, 30%, …, 80%). The threshold used was -log10(p) 3.4 that was derived from 1000x permutation at an FDR 0.05. The number of variance explained, fraction of false QTL, detected QTL, undetected QTL, as well as QTL effect size estimation were given.

**Supplementary Table S6**. QTL table of the body-size traits across different temperatures and developmental stages. Per trait in temperature and developmental stage combination, the QTL detection was done by calculating the -log_10_(p) score and significance was determined if the QTL -log_10_(p) value is bigger than the FDR (0.05) at -log_10_(p) of 3.4 based on permutation analysis. Per QTL in developmental stage and temperature combination, the position of QTL in chromosome, QTL position in base pairs, QTL left region, QTL right region, QTL effect, and the R^2^ value of each QTL are given.

**Supplementary Table S7**. Calculated broad sense and narrow sense heritability of plasticity. Plasticity per trait, developmental stage and temperature ranges the broad-sense heritability *H^2^* was calculated (H2_REML), the value for the 0.05 FDR threshold is also given (H2_FDR) as is whether the *H^2^* was significant (H2_significance). Similar for the narrow-sense heritability *h^2^* was calculated (h2_REML), the value for the 0.05 FDR threshold is also given (h2_FDR) as is whether the *h^2^* was significant (h2_significance).

**Supplementary Table 8**. Plasticity QTL table of the body-size traits across different temperatures changes and developmental stages. Per trait and developmental stage, the temperature change, the position of QTL in chromosome, QTL position in base pairs, QTL left region, QTL right region, QTL effect, and the R^2^ value of each QTL are given. The QTL detection was done by calculating the -log_10_(p) score and significance was determined if the QTL -log_10_(p) value is bigger than the FDR (0.05) at -log_10_(p) of 3.4 or 4.1 (for temperature range 24°C to 26°C).

**Supplementary Figure S1**. (A) Genetic map of the 40 RILs population. The x-axis indicates the genomic position with separate chromosomes indicated at the top of the graph. The strains of the RILs are depicted n y-axis. The RILs were genotyped by PCR-based genotyping. Orange color represents N2 genotype while CB4856 is represented by blue. Regions with uncertain genotyped are indicated by white. (B) Genotype distribution of the 729 markers along the six chromosomes. Orange represent N2 background while CB4856 was depicted in blue. The proportion of alleles frequencies that formed the RILs genetic background were roughly equal over most of the genome (Figure S2). On the left arm of chromosome I however, there was a skew of allele frequencies which we have been expected as we used the same RILs panel developed by (Li *et al*., 2006). This skew is the result of the genetic incompatibility between *zeel-1* and *peel-1* allele from N2 parent. The strains that are not protected by N2-provided *zeel-1* gene would be killed by the toxic effect from the N2-provided *peel-1* gene (Seidel *et al*., 2011).

**Supplementary Figure S2**. Correlation plot between markers. The markers were arranged to conform their physical position starting from telomere until the arms of chromosome. The correlation ranges from -1 (purple) to 1 (green) where strong correlation indicated a linkage between markers. The genotyping was done using 729 SNPs marker generated previously from low coverage sequencing of the parental strains (N2 and CB4856) (Thompson et al., 2015). To validate the genetic map for QTL mapping, we conducted a correlation analysis of the markers to detect linkage over the map. The markers are arranged to represent their physical position in chromosome.

**Supplementary Figure S3.** The reaction norms of *C. elegans* body-size traits plasticity across different temperature and developmental stages. The x-axis represents temperatures used while y-axis represent the mean value of the individual strains in their respective traits (in mm, except for body volume in mm3 and surface area in mm2). Please note that the y-axis is not identical between traits. Both parents are depicted in blue (CB4856) and orange (N2), while the RILs are grey. Lines are the traits mean value (N = 3 for RILs, and N= 9 to 12 for parents).

**Supplementary Figure S4.** Correlation analysis of body-size traits in the RIL population. (A) correlation heatmap of body-size traits of all RILs in L4 stage. There were three five strong positively correlated clusters which were temperature specific. (B) correlation heatmap of body-size traits of all RILs in adult stage. There were five strong cluster of positive correlation of worm grown in 20°C and 24°C.

**Supplementary Figure S5**. Principal component analysis of *C. elegans* body-size at (A) L4 stage and (B) adult stage: the first principal component (PC1) versus the second principal component (PC2) of the body-size traits from four different temperatures (16°C, 20°C, 24°C, and 26°C) and genetic backgrounds (N2, CB4856, and RILs).

**Supplementary Figure S6**. The body size parameters of *C. elegans* across 40 RILs and two parental strains. The average traits are expressed as percentage of sum of each traits (x-axis), followed by the mean value of the traits (y-axis). Please note that the y-axis is not identical between traits. The 40 RILs are indicated with grey, the parental N2 in orange, and the parental CB4856 in blue. Points are the traits mean value (N = 3 for RILs, and N= 9 to 12 for parents).

**Supplementary Figure S7**. Description of genotype at the peak marker locations of the signficant QTL. X-axis represents parental genotype at the peak marker location of the QTL while y-axis represents the corresponding body-size traits. Blue box visualizes CB4856 and orange bos is N2. R^2^ shows how much the variation of the body-size traits can be expalined by genetic variation in the chromosome.

**Supplementary Figure S8**. QTL plot of the body-size traits. The QTL analysis were performed across four temperatures (16°C, 20°C, 24°C, 26°C) and two developmental stages (L4 and adult). X-axis displays genomic position in the chromosome corresponding to the box above the line while y-axis represents the LOD score. Box in the right graph show the developmental stages. Red line represents QTL at 16°C, green line at 20°C, blue line at 24°C, and purple line at 26°C. Red-dash line represents LOD threshold (1000x permutation-based significant LOD with FDR = 0.05).

