## Supplementary figures and images for "The genetic architecture underlying body-size traits plasticity over different temperatures and developmental stages in *Caenorhabditis elegans*"

### Figure S1

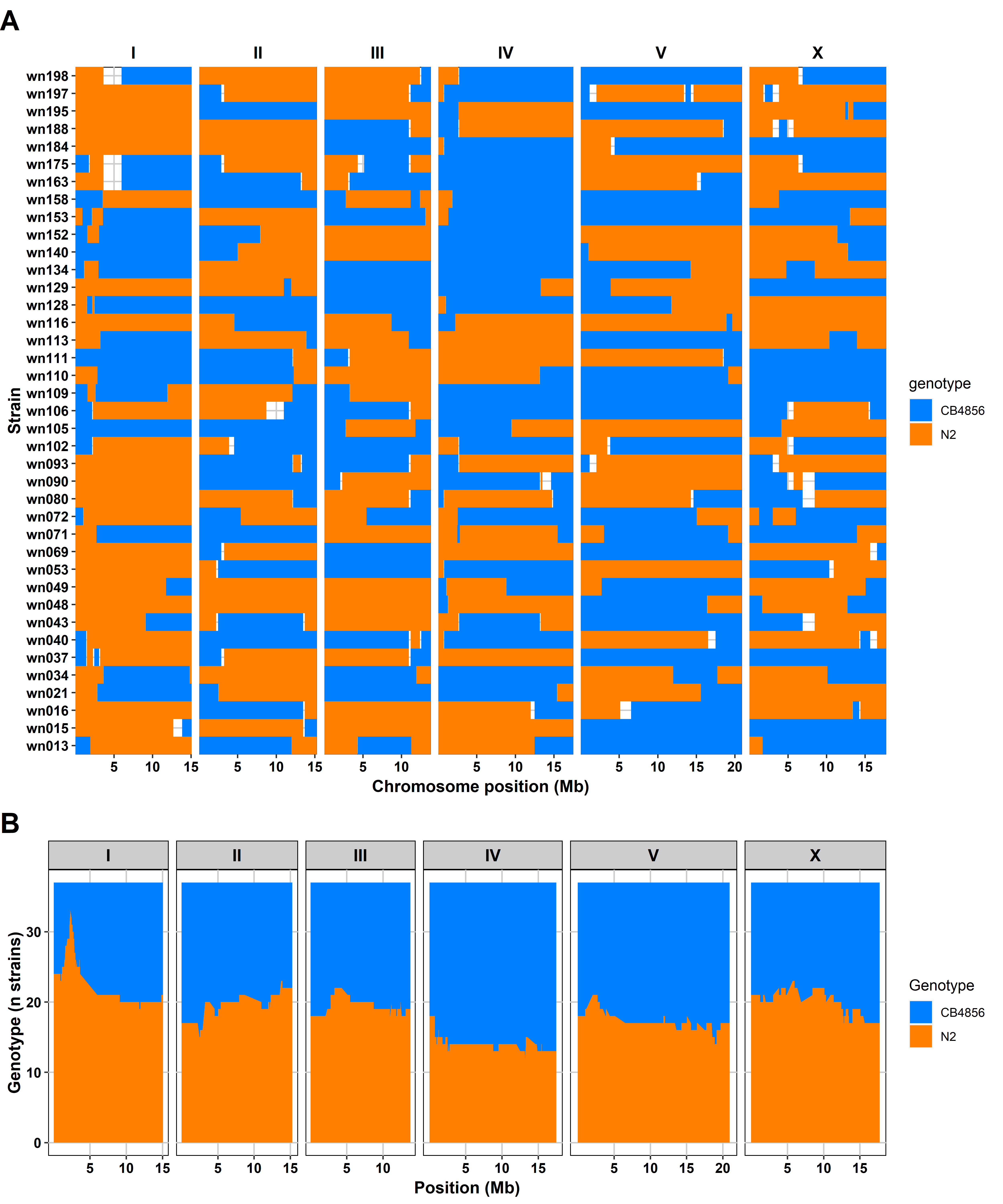

### Figure S2

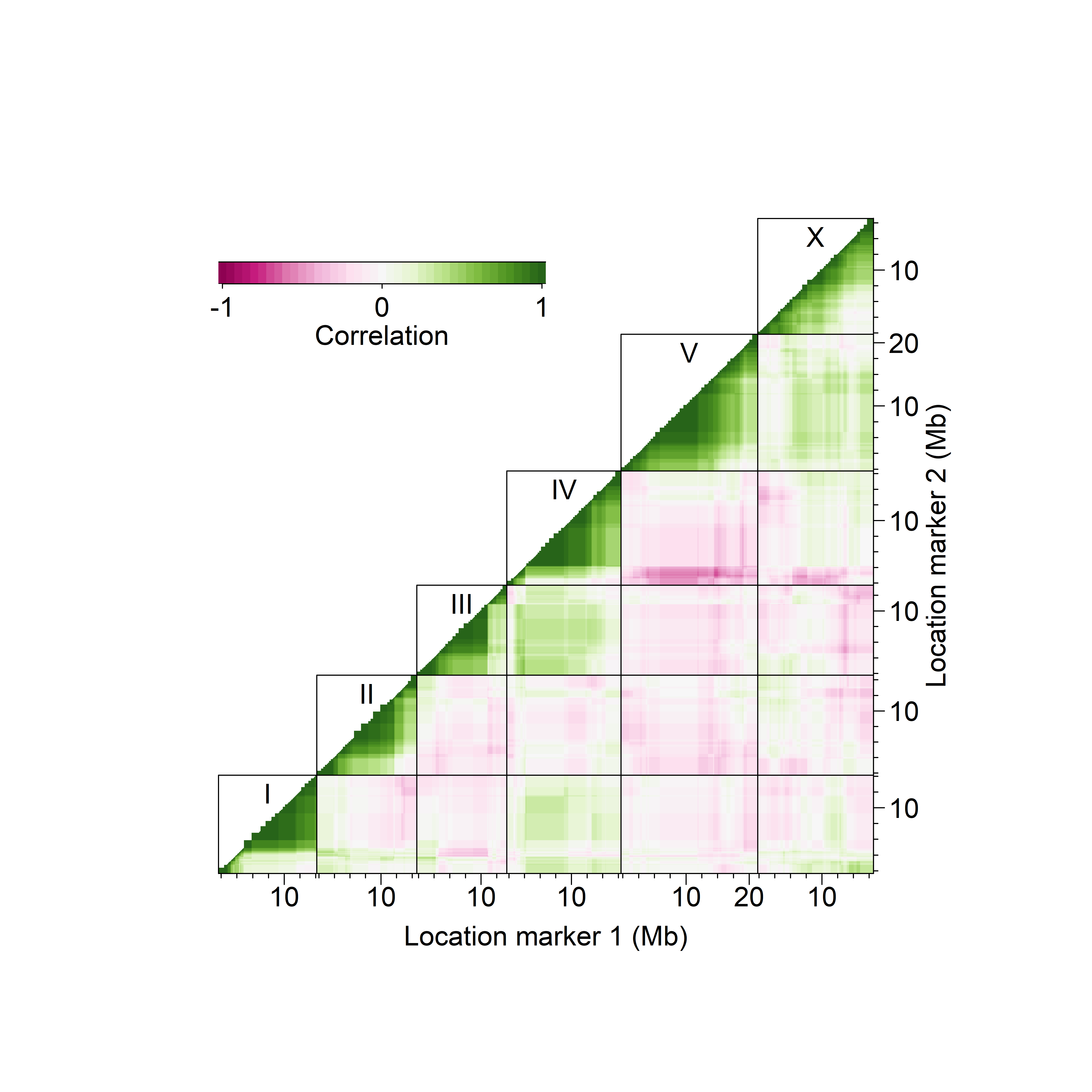

### Figure S3

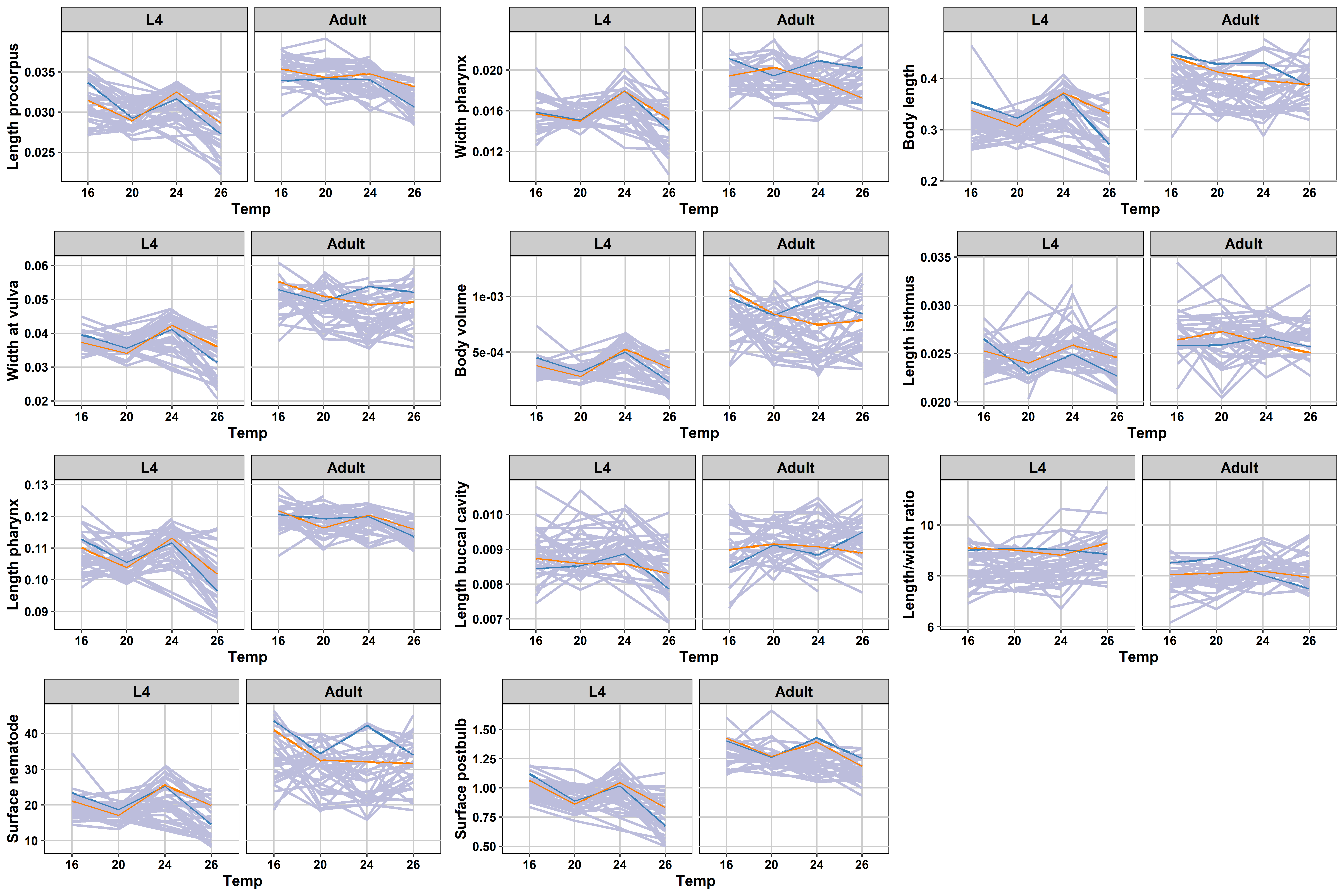

### Figure S4A

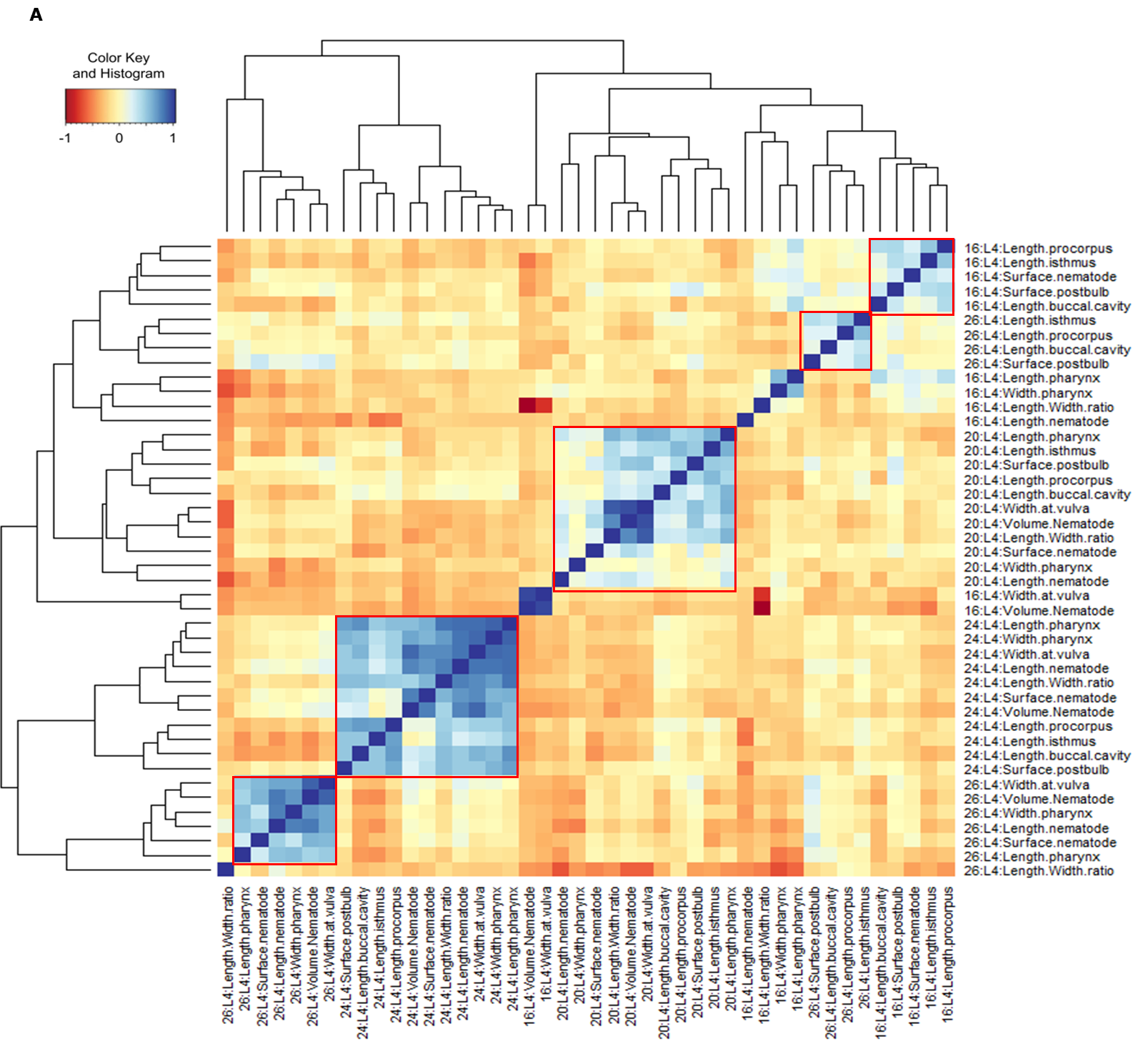

### Figure S4B

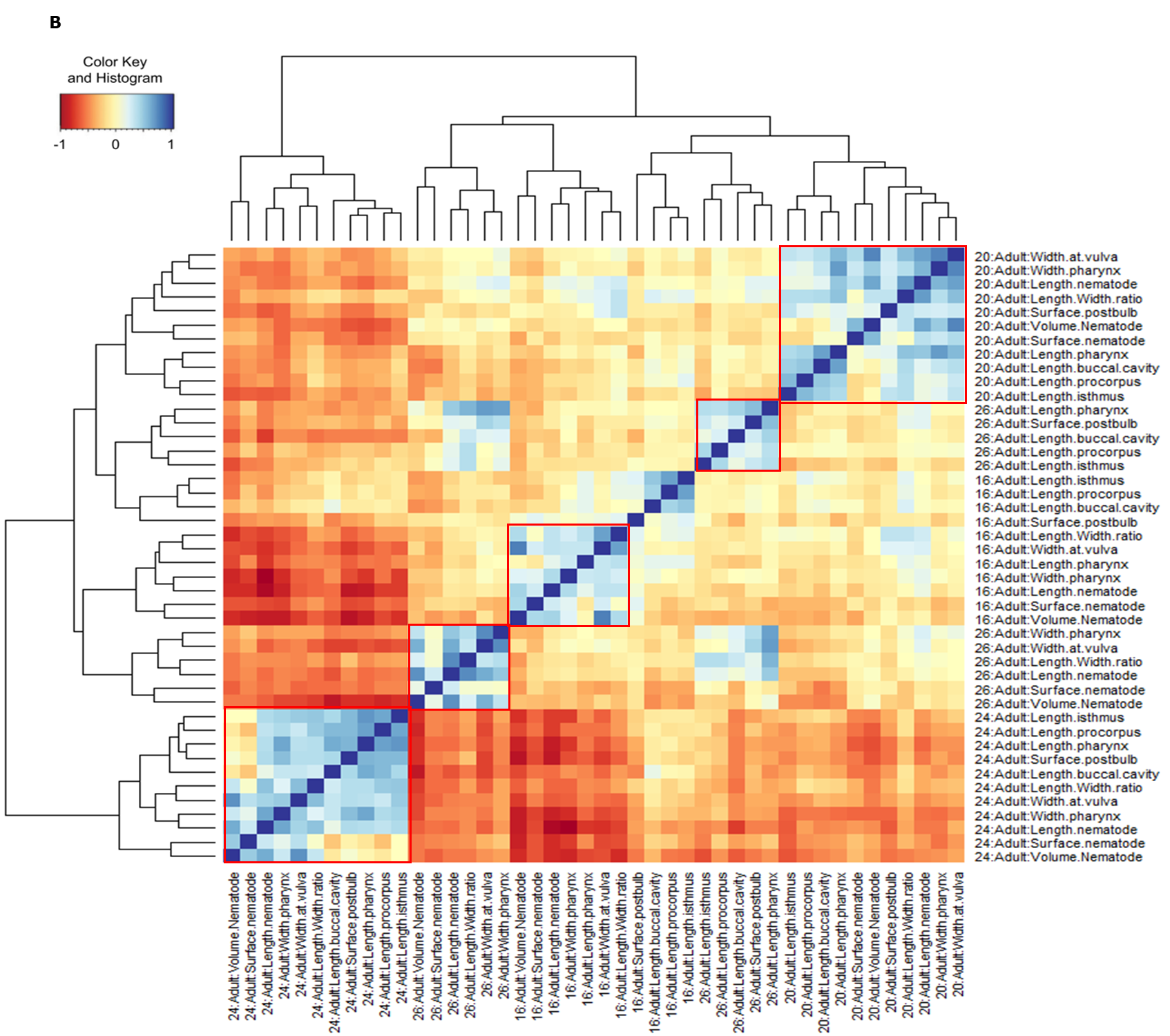

### Figure S5

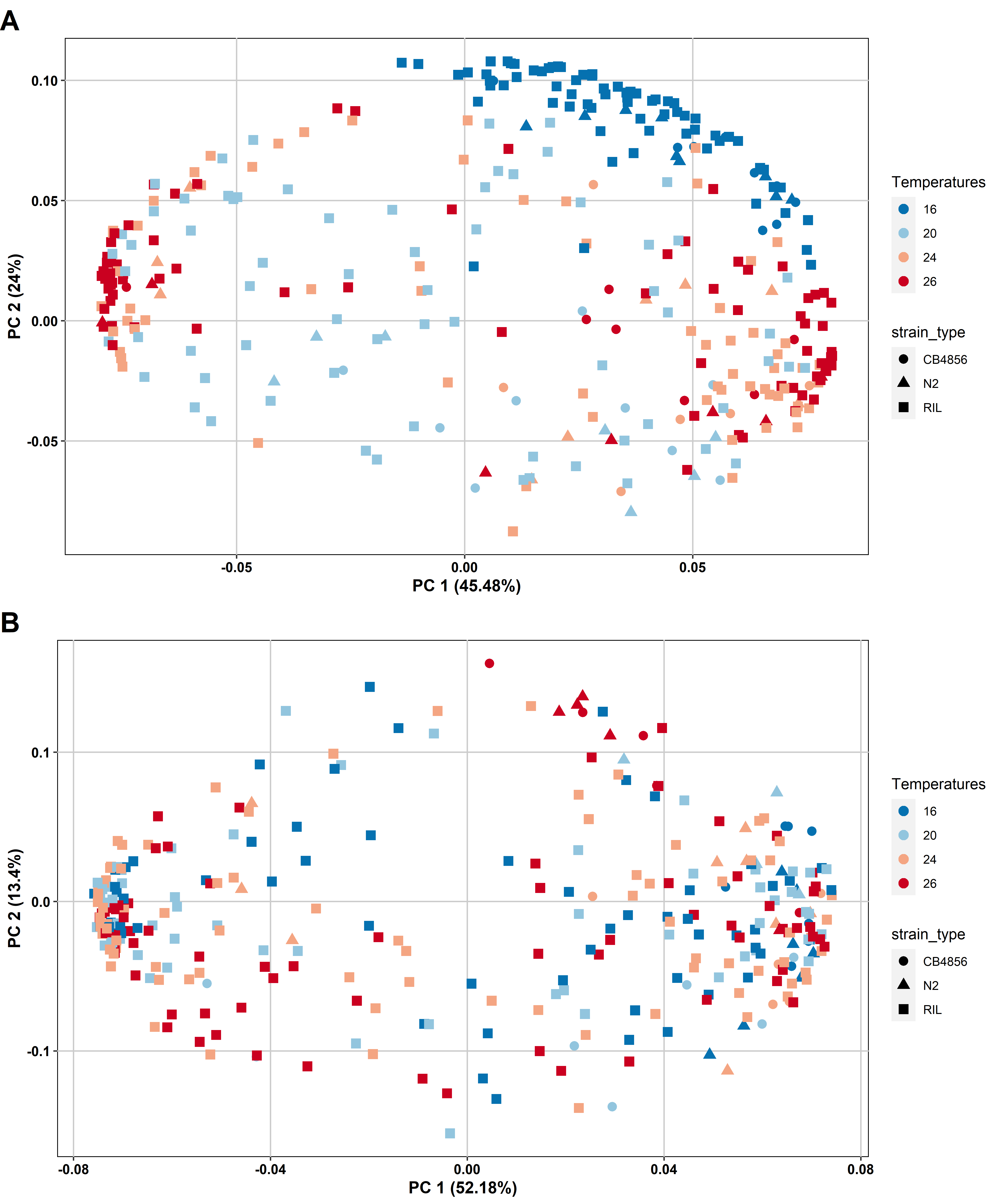

### Figure S6

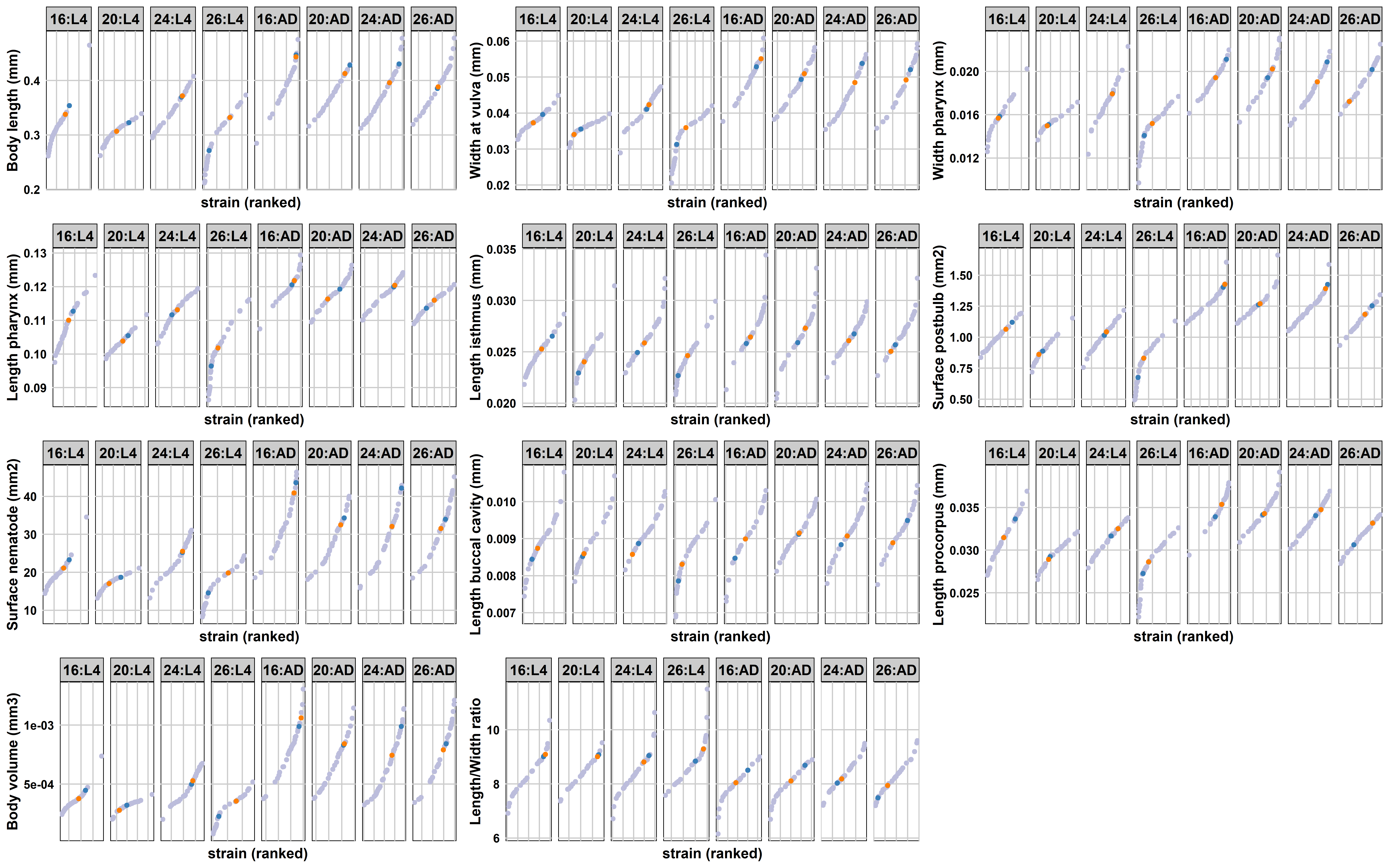

### Figure S7

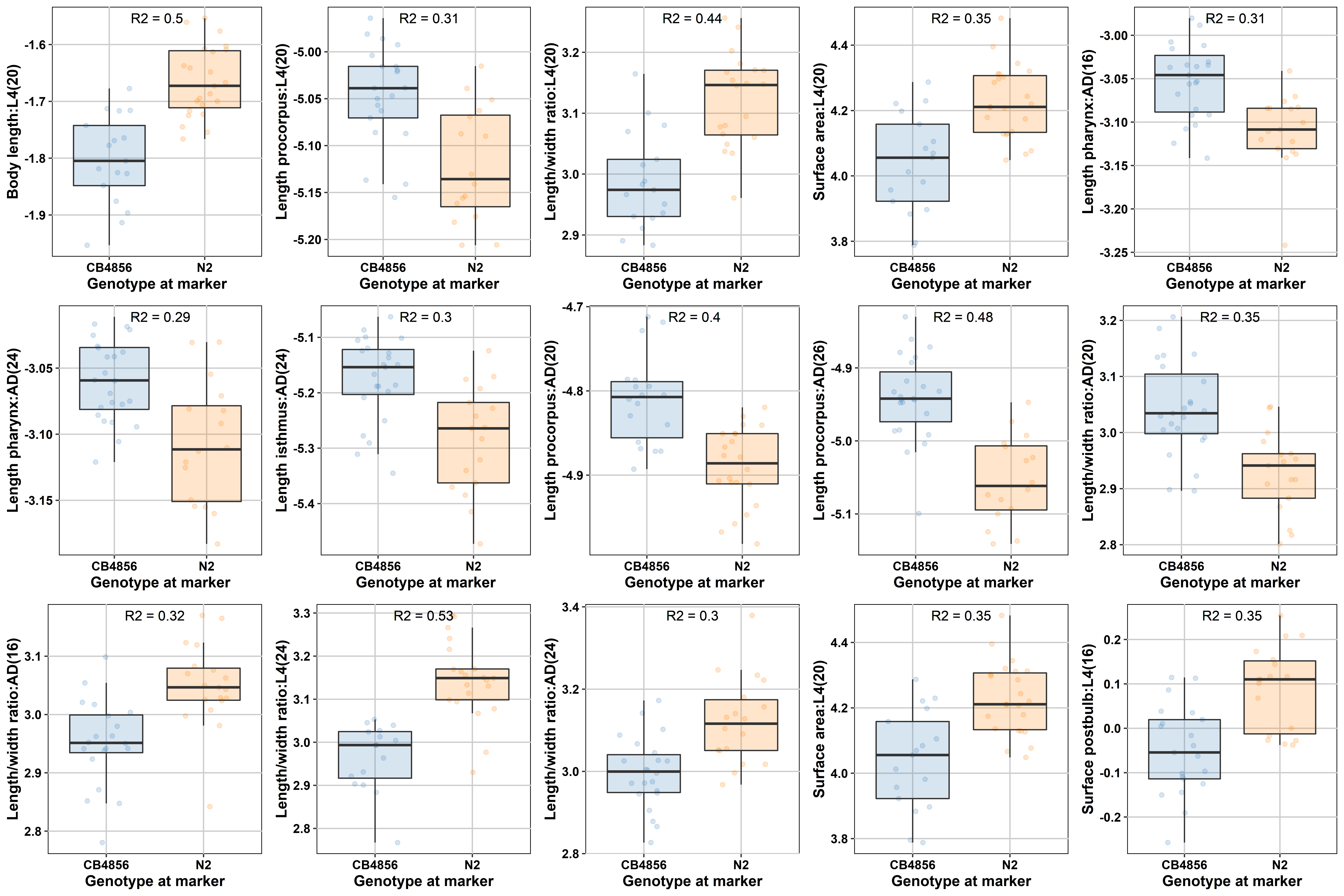

### Figure S8

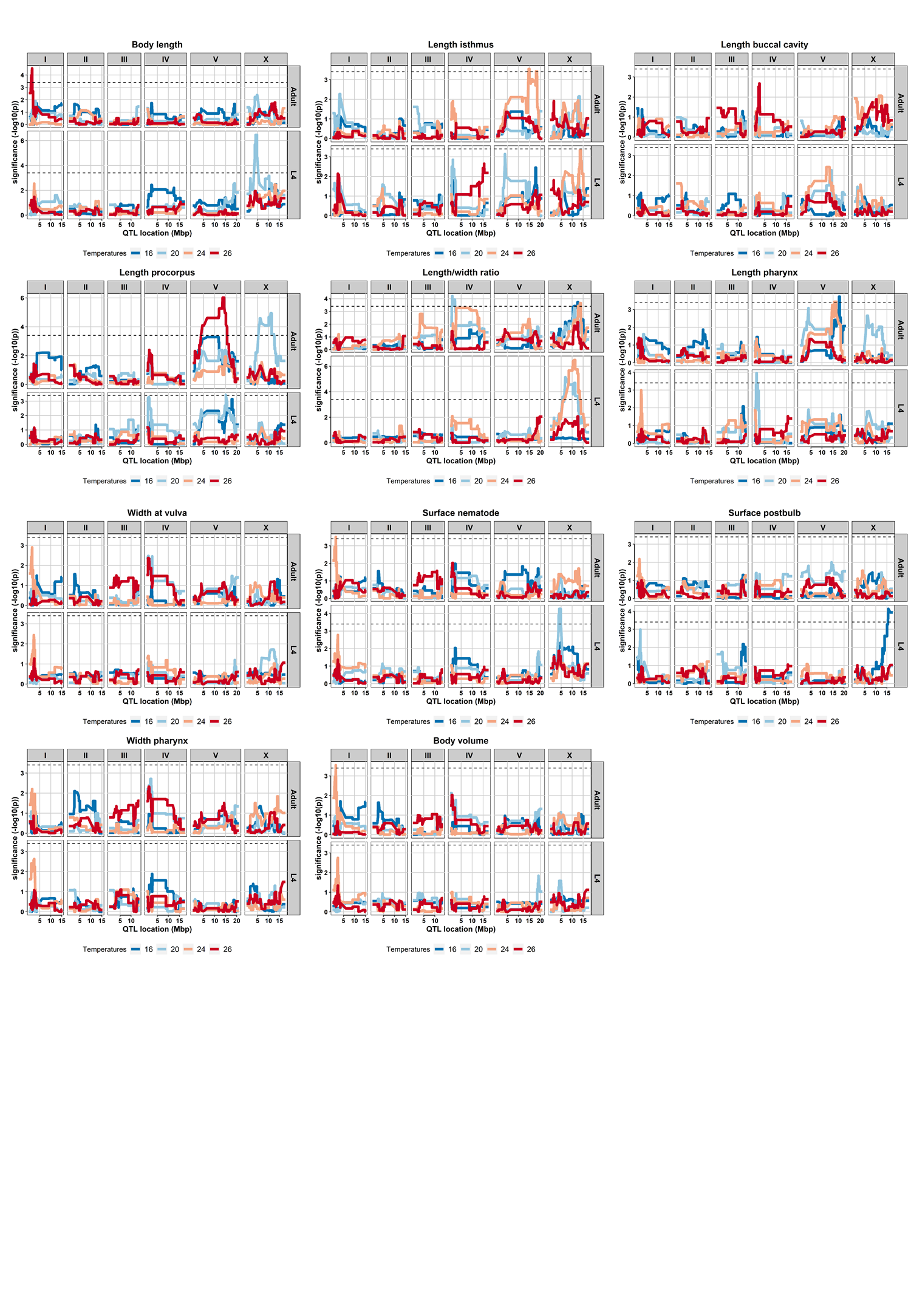
